# Dissociating the impact of movement time and energy costs on decision-making and action initiation in humans

**DOI:** 10.1101/2021.05.26.445778

**Authors:** Clara Saleri Lunazzi, Amélie J. Reynaud, David Thura

**Affiliations:** Lyon Neuroscience Research Center – ImpAct team, Inserm U1028 – CNRS UMR5292 – Lyon 1 University, 16 avenue du Doyen Lépine, 69675 Bron - France

## Abstract

Recent theories and data suggest that adapted behavior involves economic computations during which multiple trade-offs between reward value, accuracy requirement, energy expenditure and elapsing time are solved so as to obtain rewards as soon as possible while spending the least possible amount of energy. However, the relative impact of movement energy and duration costs on perceptual decision-making and movement initiation is poorly understood. Here, we tested 31 healthy subjects on a perceptual decision-making task in which they executed reaching movements to report probabilistic choices. In three distinct blocks of trials, the reaching time and energy costs were independently varied while decision difficulty was maintained similar at the block level. Participants also performed a fully instructed delayed-reaching (DR) task in each motor condition. Results in that DR task show that time-consuming movements extended reaction times (RTs) in most subjects, whereas energy-consuming movements led to mixed effects on RTs. In the choice task, about half of the subjects decreased their decision durations (DDs) in the time consuming condition, while the impact of energy costs on DDs were again mixed across subjects. Decision accuracy was overall similar across motor conditions. These results indicate that movement duration and, to a lesser extent, energy expenditure, idiosyncratically affect perceptual decision-making and action initiation. We propose that subjects who shortened their decisions in the time consuming condition of the choice task did so to limit a drop of their rate of reward.

## INTRODUCTION

For humans and animals in general, life presents a constant stream of decisions about actions to make regarding food, mobility, social interactions and many other situations. Crucially, decision and action often involve context-dependent computations during which effort is traded against time to obtain rewards as soon as possible while spending the least possible amount of energy (Shadmehr and Ahmed 2020). These computations are complex to solve because strong interactions exist between reward valuation, elapsing time and energy expenditure (Figure 1A). For example, human and non-human primates expecting large rewards reduce their reaction time and increase the vigor of the movements executed to obtain these rewards (Kawagoe et al. 1998; Manohar et al. 2015; Reppert et al. 2015; Revol et al. 2019; Yoon et al. 2018). But increasing vigor usually means increasing energetic expenditure, which discounts reward value (Klein-Flügge et al. 2015; Sugiwaka and Okouchi 2004). Indeed, when an effortful movement is anticipated, reaction times are increased, and action vigor is reduced (Morel et al. 2017; Summerside et al. 2018). Besides, to be rewarded, it is often necessary to provide accurate movements. A fundamental and long-established observation is the so-called speed-accuracy trade-off: when actions are performed faster, they tend to be less precise (Fitts 1954). This principle applies to both motor and cognitive performances (Heitz 2014). Individuals could thus benefit from maximizing accuracy and minimizing effort by making slow movements. However, this strategy implies increasing behavior duration, which inevitably delays the completion of the task and the acquisition of the reward, leading to the well-known temporal discounting of reward value (Berret and Jean 2016; Choi et al. 2014; Haith et al. 2012; Myerson and Green 1995; Shadmehr et al. 2010). To summarize, both time and effort discount the value of reward, and reducing reward temporal discounting requires increasing energy expenditure, which in turn discounts the value of reward too.

**Figure 1:**
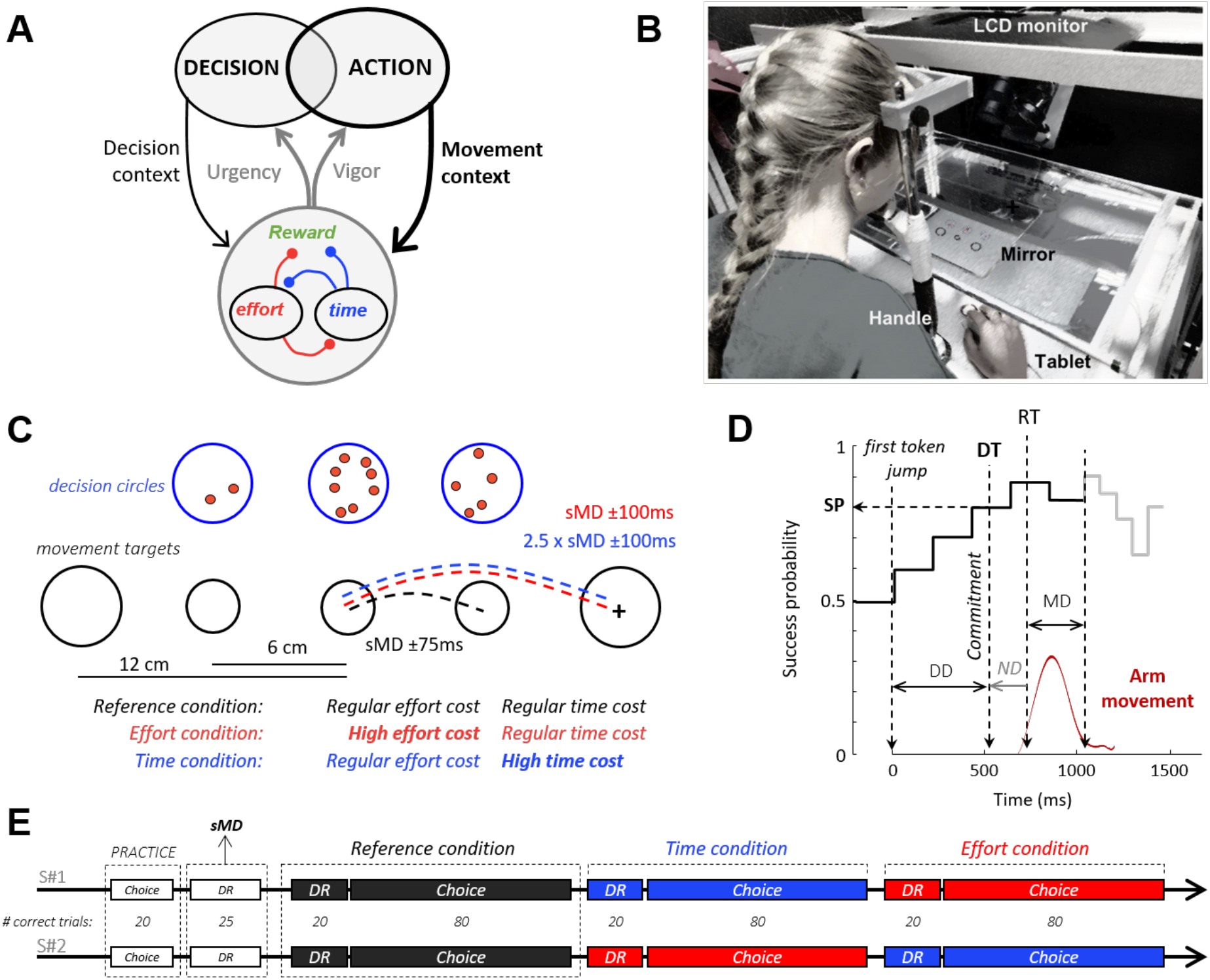
Theoretical framework, experimental set-up and design. **A**. To maximize the rate of reward during goal-directed behavior, decision and action must be coordinated. Regulation signals (gray arrows) allow such coordination. These signals are determined based on the decisional (thin black arrow) and motor (bold black arrow) context-dependent integration of reward value, elapsed time and energy expenditure. These three components are intertwined in a trade-off of reciprocal negative interactions (red and blue connections). **B**. Experimental apparatus (see text for details). **C**. Visual display and motor conditions in the choice task. Blue circles illustrate the decision stimuli. Tokens successively jump from the central circle to one of the two lateral circles. Black circles show the movement stimuli. Subjects move a handle (cross) from a central start circle to one of the two lateral targets, depending on their choice. Movement amplitude and duration (MD) are imposed in distinct blocks of trials. In the Reference condition (black), both lateral movement targets are located close to the starting circle and a specific movement duration (with a 150ms tolerance window) is imposed. In the two other conditions, both lateral movement targets are located twice as far apart from the starting circle compared to the Reference condition. In the Effort condition (red), the imposed movement duration is the same as in the Reference condition whereas in the Time condition (blue), the imposed movement duration is about twice longer. For these two costly conditions, the tolerance interval around the imposed movement duration is 200ms. **D**. Temporal profile of success probability in one example trial of the choice task. At the beginning of the trial, each target has the same success probability (0.5). When the first token jumps into one of the two potential targets (the most leftward vertical dotted line), success probability of that target increases to ∼0.6. Success probability then evolves with every jump. Subjects execute a reaching movement (red trace) to report their choices. Movement onset (RT) and offset times are used to compute movement duration (MD), and movement offset marks the moment when the tokens that remain in the central decision circle jump more quickly to their assigned target (gray trace). The estimated time of the decision (DT) is computed by subtracting the subject’s mean non-decision delay (ND) estimated in a simple delayed-reach (DR) task from movement onset time, allowing computation of the success probability (SP) at that moment. Only 10 out of 15 jumps are illustrated on this SP profile. **E**. Time course of the two sessions (#1 and #2). Subjects start each session with 25 trials of the choice task to familiarize themselves with the set-up. Then 25 trials of the DR task with no constraint on movement duration are performed in order to determine for each subject the average spontaneous arm movement duration (sMD) which will be necessary to determine the time constraints of the response movements in each condition. Subjects next need to complete 20 correct trials in the DR task and 80 correct trials in the choice task for each motor condition. They start with the Reference condition, followed by the Time and the Effort conditions in the first session. The order of presentation of the two costly conditions is reversed in the second session.

What are the implications of these relationships during goal-directed behavior? For anyone making a decision, the most adaptive strategy is to choose options that maximize the global rate of reward (Balci et al. 2011; Bogacz et al. 2010), which occurs when *both* decision and action are sufficiently accurate but not overly effortful and time consuming. Because trade-offs during decision and action have been typically studied in isolation, mechanisms allowing a coordinated computation of reward rate are still elusive. Recent promising advances suggest however that motor control and choices are governed by partly overlapping principles (Carland et al. 2019; Morel et al. 2017; Yoon et al. 2018). First, motor costs influence human decision-making when choices rely on action properties only (Cos et al. 2011, 2014; Michalski et al. 2020; Morel et al. 2017) or when they are driven by perceptual stimuli (Hagura et al. 2017; Marcos et al. 2015). During motor decisions, movement duration is perceived by humans as the most effortful cost (Michalski et al. 2020; Morel et al. 2017), but whether or not this result generalizes beyond motor choices is unknown. Second, in the foraging paradigm where one makes decisions regarding how long to stay and collect rewards from one patch, and then move with certain speed to another patch, the goods collection duration and the vigor with which human subjects move from one reward site to another are governed by a mechanism allowing to maximize the overall capture rate (Yoon et al. 2018). Finally, both human and monkey level of decision urgency at commitment time strongly predicts the duration of the movements executed to express these choices and allows reward rate maximization (Thura 2020; Thura et al. 2012, 2014), in agreement with theoretical and experimental work (Churchland et al. 2008; Ditterich 2006; Drugowitsch et al. 2012).

Together, these studies suggest that movement vigor is coordinated with decision-making urgency to optimize the rate of reward. Recently, we provided strong support for this hypothesis by showing that when movement accuracy requirements are relaxed, decision duration is extended, allowing human subjects to increase their choice accuracy (Reynaud et al. 2020). The present work is designed to investigate this coordination between decision and action further and assess whether, how and why motor time and/or energy costs influence perceptual choices in human subjects. More specifically, we aimed at addressing three questions: (1) Does the motor context, costly or not, in which a choice is made influence decision-making? Because decision and action need to be coordinated in order to maximize the rate of reward, we predict that decision speed and/or accuracy will be modulated depending on the cost of the movement executed to express that choice; (2) What is the most impactful motor cost (time or energy) context during perceptual decision-making? Because several studies have shown that movement duration is the parameter that subjects tend to control to increase their rate of reward (Choi et al. 2014; Haith et al. 2012; Shadmehr et al. 2010), we predict that movement duration should have the largest impact on subjects’ choices; (3) How variable are these effects between subjects? Recent studies revealed individual “traits” of decision and motor behavior, showing that despite facing identical trials, some subjects could decide and act much faster than others (Berret et al. 2018; Labaune et al. 2020; Reppert et al. 2018; Thura 2020). This suggests variable sensitivities to motor costs at the population level. We thus predict that the impact of the movement energy and/or temporal costs on decision-making will be idiosyncratic.

## MATERIAL AND METHODS

### Power analysis

We performed an *a priori* power analysis to estimate the optimal combination of trials per condition and participant numbers, depending on expected effect sizes and variabilities (Baker et al. 2020). Based on a previous experiment conducted on 20 human subjects performing a similar task (Reynaud et al. 2020), we estimated a mean difference of decision duration between motor conditions of 150ms, a within-subject standard deviation (SD) of 420ms, a between-subject SD of 230ms, and we set the alpha level to 0.05. For a standard power of 80%, 22 subjects had to be tested on 32 trials per condition. To increase the power and reach 90%, we needed to test at least 28 subjects in about 80 trials per conditions. Given our past experience with similar experiments (Reynaud et al. 2020; Thura 2020), executing a minimum of 80 trials per condition in an experiment that is designed to include 3 conditions takes about one hour, which is an acceptable duration for a healthy, young subject. But to increase the statistical power of our results further without increasing session duration, we tested each participant twice, in two separate sessions. The effect of the session on the impact of motor costs on goal-directed behavior will be addressed in another publication.

### Participants

Thirty-one healthy human subjects (ages: 18-36; 20 females; 29 right-handed) participated in this study. All gave their consent before starting the experiment. The INSERM ethics committee (IRB00003888) approved the protocol on March 19th, 2019. Each participant was asked to perform two experimental sessions (with a maximum of 7 days between sessions) and they received a monetary compensation (15 euros per completed session) for participating in this study. All subjects completed the two sessions and have been included in the present dataset.

### Experimental set up

Subjects sat in an armchair and made planar reaching movements using a handle held in their dominant hand. A digitizing tablet (GTCO CalComp) continuously recorded the handle horizontal and vertical positions (100Hz with 0.013cm accuracy). Target stimuli and cursor feedback were projected by a DELL P2219H LCD monitor (60Hz refresh rate) onto a half-silvered mirror suspended 26cm above and parallel to the digitizer plane, creating the illusion that targets floated on the plane of the tablet (Figure 1B).

### Tasks and experimental design

Subjects were instructed to perform alternations of two tasks: a choice task, modified from Cisek et al. (2009), and a delayed-reaching (DR) task.

In the choice task (Figure 1C), participants faced a visual display consisting of three blue circles (the decision circles; 1.5cm radius) placed horizontally at a distance of 6 cm of each other and three black circles positioned 12cm below (the movement targets). At the beginning of each trial, 15 red tokens are randomly arranged in the central blue circle. The position of the decision stimuli was constant, but the distance between the central and lateral movement targets varied, set to either 6cm (short distance) or 12cm (long distance) from the central circle in distinct blocks of trials. The size of the central movement circle (the starting circle) was constant (0.75cm radius) whereas the size of lateral movement circles was set to either 1cm radius in the short distance trials or 1.5cm radius in the long distance ones to minimize the perceived size and accuracy requirement differences between conditions (Sperandio and Chouinard 2015). Importantly, the size of the movement target was chosen to be large enough to minimize the motor accuracy constraints and avoid major speed-accuracy tradeoff adjustments. The effect of motor accuracy on decision-making has been investigated in a recent publication (Reynaud et al. 2020).

A choice task trial (Figure 1D) starts when the subject places the handle in the starting position and remains still for 500ms. The tokens then start to jump, one by one, every 200ms, in one of the two possible lateral blue circles. The subject has to decide which of the two decision circles will receive the majority of the tokens at the end of the trial. To report a decision, the subject has to move and hold the handle into the lateral movement target corresponding to the side of the chosen decision circle for 500ms. Subjects were allowed to make and report their choice at any time between the first and the last token jump. The tokens that remain in the central circle once the target is reached then jump more quickly (every 50ms), motivating subjects to answer before all tokens have jumped to increase their rate of correct decisions. Note that this feature entails that movement duration carries a temporal cost with respect to the subject’s rate of correct decisions. A visual feedback about decision success or failure (the chosen decision circle turning either green or red, respectively) is provided after the last jump. A 1500ms period (the inter-trial interval, ITI) precedes the following trial.

The delayed-reach (DR) task is similar to the choice task except that only one lateral decision circle along with its associated movement target are displayed at the beginning of the trial (either at the right or at the left side of the central circle with 50% probability). Moreover, all tokens move from the central circle to this unique circle at a GO signal occurring after a variable delay (1000 ± 150ms). This fully instructed task is used to estimate the spontaneous movement duration of each subject and their mean reaction (i.e. non-decision) time in each motor condition (see below).

At the beginning of the session, a practice period consisting of performing 20 choice task trials with short and long distance targets (with 50% probability) was proposed, mainly allowing subjects to get familiar and comfortable with the manipulation of the handle on the tablet. Then, the subject had to perform 25 trials of the DR task with short distance movements and no constraint on movement duration. This block of trials was used to determine for each subject the average spontaneous arm movement duration (sMD) necessary to reach short distance targets. Based on this duration, we determined for each subject the spontaneous MD interval (sMD ± 75ms) and the long-distance MD interval (2.5 x sMD ± 100ms). The first six subjects performed the tasks with a lower temporal tolerance (± 50ms and ± 75ms for the short and long distance movements, respectively), but based on their motor performance and post-session interviews, we decided to relax the temporal constraints of the movements for the rest of the population (± 75ms and ± 100ms for the short and long distance movements, respectively). Subjects then performed alternations of DR and choice task trials in the three different motor conditions described in the next paragraph (Figure 1E).

To assess the influence of the time and energy costs of movements on decision-making, the position of the lateral movement targets as well as the movement duration (MD) interval allowed to reach these targets were varied in three distinct blocks of trials (Figure 1C). In the “Reference” condition, subjects were instructed to execute short distance movements, within their spontaneous MD interval. In the “Time” condition, subjects had to execute long distance movements within their long MD interval, thus doubling movement duration (Figure 2A, left) without much of an increase of energy expenditure compared to the Reference condition (Figure 2B). In the “Effort” condition, subjects were instructed to execute long distance movements, just as in the Time condition, but within their spontaneous MD interval. This Effort condition thus required about twice faster movements than the Reference condition, significantly increasing their energy cost (Figure 2B) without increasing their duration (Figure 2A, right). Importantly, the decision component of the task was strictly similar between the three motor conditions. For instance, the maximum decision time allowed (15 token jumps, 2800ms) wasn’t shorter in the Time condition compared to the two other blocks. A trial was considered incorrect if movement did not meet these block-dependent spatio-temporal constraints. In this case, the subject received a visual feedback (both movement targets turned red) as well as a 500ms audio feedback indicating that movement was too fast or too slow (800 Hz or 400 Hz sound, respectively). If the movement was executed within the imposed duration interval and the subject chose the target receiving the majority of tokens at the end of the trial, that trial was considered as correct. The goal for each subject was to perform in each of the two sessions 20 correct DR task trials in each condition and 80 correct choice task trials in each condition, motivating them to maximize their rate of correct responses. Subjects started both sessions in the Reference condition. In the first session, the Reference condition was followed by the Time and the Effort conditions, whereas this order was reversed in the second session.

**Figure 2:**
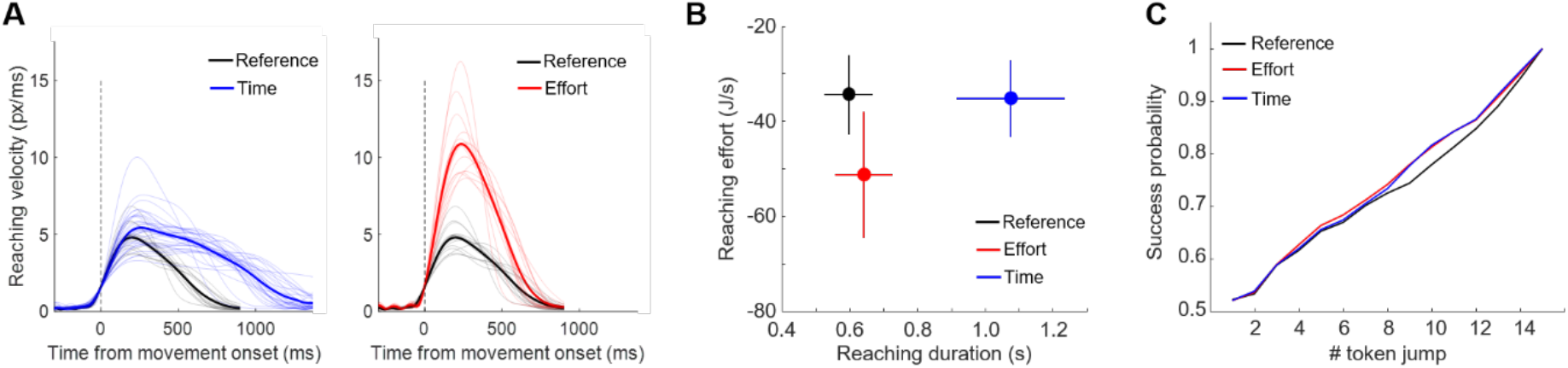
Control analyses. **A**. Individual (thin) and average (bold) reach velocity profiles in the three motor conditions of the choice task, aligned on movement onset. Only adequate movements are included. Time and Reference conditions are compared in the left panel; Effort and Reference conditions are compared in the right panel. **B**. Relationship between the average (±SD) reaching duration and effort across subjects in the three motor conditions (see eq. 2 and details in text). **C**. Average success probability profiles of trials experienced by subjects in each of the three motor conditions (eq. 3).

### Data analysis

The present analyses were performed on trials collected from all subjects performing both sessions #1 and #2. Data were analyzed off-line using custom-written MATLAB (MathWorks) and R (https://www.r-project.org/) scripts. Unless stated otherwise, data have been combined across sessions, and are reported as mean ± standard deviation (SD).

We first analyzed the kinematic properties of the reaching movements performed by subjects in each of the three motor blocks. Horizontal and vertical handle position data were filtered using a fifteen-degree polynomial filter and then differentiated to obtain velocity profiles (see one example reach velocity profile depicted in Figure 1D). Movement onset and offset were determined using a 3.75 cm/s velocity threshold. Peak velocity and movement duration (MD) were respectively computed as the maximum value and the time between these two events.

In the present work, we manipulated subjects’ reaching speed (for a given amplitude) to vary movement energetic expenditure between conditions, as reaching speed and metabolic rate strongly co-vary (Ludlow and Weyand 2016; Shadmehr et al. 2016). But we also estimated, post hoc, the energy *er* spent during each reaching movement as a function of the reaching distance *d* (in meters) and duration *t* (in seconds) using the following equation, from Shadmehr et al.’s study (2016) in which the energetic cost of 2D reaching movements was measured (via expired gas analysis) and parametrized as a function of movement duration, arm mass, and distance:

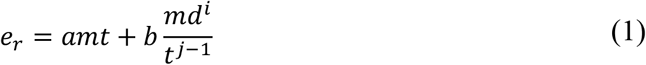

In eq. 1, *m* is a constant which represents the mass of the arm, estimated in the present work based on subjects’ weight data (m=weight *∼0.05, de Leva 1996). Terms *a, b, i* and *j* are fixed coefficients determined in Shadmehr et al.’s (2016) experiment. For the present estimations, we set a=15, b=77, i=1.1 and j=2.7. Energetic consumption may represent an objective measure of movement effort. We used this estimation to compute participants’ expected reward rate in the choice task (see eq. 4). In the context of reaching movements however, past studies proposed that effort is rather subjectively perceived as the temporally discounted metabolic cost of performing an action (Körding et al. 2004; Shadmehr et al. 2016), resulting in the following equation:

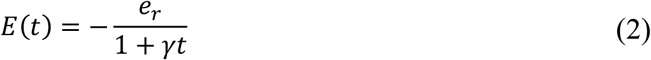

where *γ* is the hyperbolic temporal discounting parameter. Thus, assuming that movement duration delays the acquisition of reward, the act of moving fast leads to acquisition of a large reward in exchange for a large effort, whereas moving slowly leads to acquisition of smaller, discounted reward later in exchange for payment of small effort. We used this metric to control for the efficiency of our experimental conditions to dissociate reaching duration from effort (Figure 2B).

The analysis of participants’ decision-making behavior in the choice task focused on the duration of the decision (DD) and the success probability of the choice (SP). To estimate the time at which subjects committed to their choice on each trial, we first defined the reaction time (RT) as the time of movement onset with respect to the first token jump. We then subtracted from each RT the mean non-decision delay estimated based on subjects’ RTs in the same motor condition of the DR task, providing the time at which the deliberation ends (decision time, DT). RTs measured in the DR task also allowed us to assess the effects of the motor context on movement initiation (Figure 3). Then, DD was computed as the duration between the first token jump and DT in the choice task (Figure 1D).

**Figure 3:**
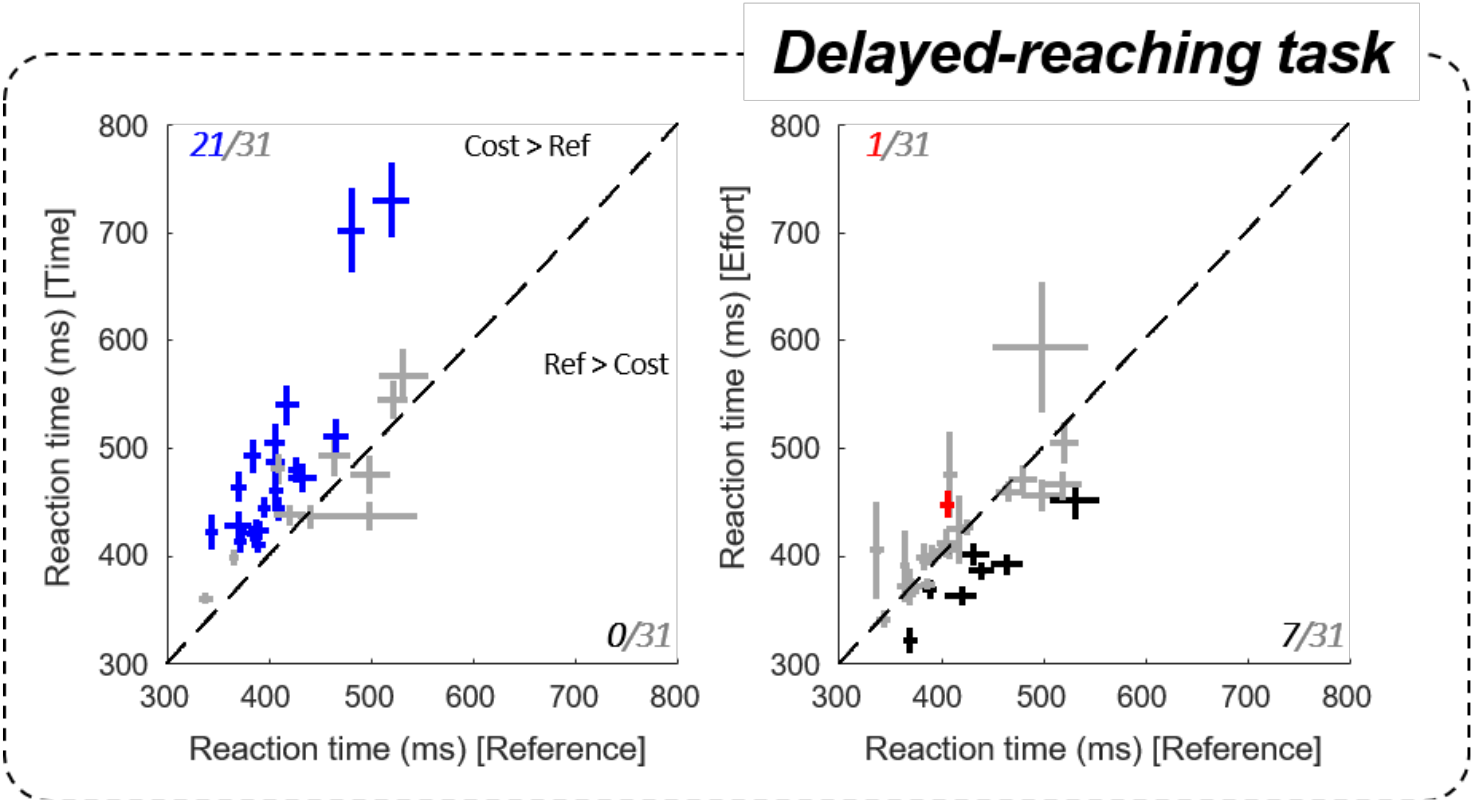
Effect of motor conditions on motor initiation duration. Comparison of subjects’ reaction times (RTs) in the delayed-reach task between the costly conditions (ordinate, left panel: Time / right: Effort) and the Reference condition (abscissa). Pluses indicate the mean and SE for each subject. Left panel: blue pluses indicate data that are significantly larger in the Time condition compared to the Reference condition. Right panel: red (black) pluses indicate RTs that are significantly larger (smaller) in the Effort condition compared to the Reference condition.

The choice task design allows to calculate, at each moment in time during a trial, the success probability *pi(t)* associated with choosing each target *i* (eq. 3). For instance, for a total of 15 tokens, if at a particular moment in time the right target contains NR tokens, whereas the left contains NL tokens, and there are NC tokens remaining in the center, then the probability that the target on the right will ultimately be the correct one, i.e. the success probability (SP) of guessing right is as follows:

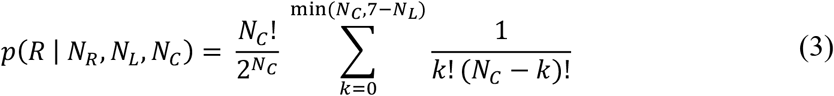

To control for the average decision difficulty between motor conditions and make sure that this difficulty could not account for potential differences in the subjects’ decision strategy, all subjects faced the same sequence of trials in which we interspersed among fully random trials (20% of the trials in which each token is 50% likely to jump into the right or into the left lateral circle) three special types of trials characterized by particular temporal profiles of success probability. Subjects were not told about the existence of these trials. 30% of trials were so-called “easy” trials, in which tokens tended to move consistently toward one of the circles, quickly driving the success probability *pi(t)* for each toward either 0 or 1. Another 30% of trials were “ambiguous”, in which the initial token movements were balanced, making the *pi(t)* function close to 0.5 until late in the trial. The last special trial type was called “misleading” trials (20 %) in which the 2-3 first tokens jumped into the incorrect circle and the remaining ones into the correct circle. Crucially, the sequence was designed so that the proportion of trial types was continuously controlled and kept constant within and between motor conditions. Even if the above criteria leave some room for variability within each trial type, the sequence provided subjects with an overall same level of decision difficulty between motor conditions (Figure 2C). In all cases, even when the temporal profile of success probability of a trial was predesigned, the actual correct target was randomly selected on each trial.

We computed subjects’ mean reward rate in each motor condition in order to assess the impact of each motor cost on this metric. We quantified the expected reward rate in each trial *i* with the following equation:

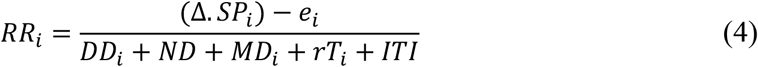

where Δ is a hypothetic value (in Joules) assigned by the brain to a positive outcome in each trial, SP is the probability of choosing the correct target in trial *i* (eq. 3), *e* is the energetic consumption of reaching toward the chosen target in trial *i* (eq. 1), DD is the decision duration for trial *i*, ND is the condition-dependent non-decision delay (estimated in the DR task), MD is the duration of the movement in trial *i*, rT is the time taken by the remaining tokens to jump in their assigned target in trial *i*, and ITI is the fixed inter-trial interval. We then computed the average reward rate across trials and compared this average rate between motor conditions at the population level.

### Statistics

We used linear mixed effects models to examine the effect of motor conditions on the different dependent measures described above for each subject. Analyses were performed using the ‘lme4’ package for R (Bates et al. 2015). We defined a model containing the most appropriate random effects (i.e. factors of non-interest) for each variable using Likelihood Ratio Tests. For the DR task variable (i.e. RT), *sessions* (#1 and #2) were included in the model as random intercept. For the choice task motor (duration, velocity peak, effort) and decision (duration, success probability and reward rate) variables, *sessions* (#1 and #2) and *trial types* (random, easy, misleading and ambiguous) were included as a random intercept. We then tested the effect of the motor conditions (Reference, Time and Effort) as a fixed factor in order to evaluate their influence on each dependent variable tested. We also computed a linear mixed effect model for each dependent variable tested across all subjects by adding *subjects* as a random factor. Finally, post-hoc comparisons were carried out using pairwise comparisons through the ‘lsmeans’ package for R (p-adjusted with false discovery rate method, Benjamini and Hochberg 1995; Lenth 2016) to assess the effect of the different motor conditions (Reference vs. Time and Reference vs. Effort).

## RESULTS

In the choice task, the objective in a given session was to perform 80 correct trials in each motor condition. A trial was considered correct when the subject both chose the correct target and executed an appropriate movement to report it. A movement was appropriate when it stopped in the target within the determined time interval. Across the two sessions, subjects performed 179 ± 36 trials (average ± SD, correct and incorrect) in the Reference condition, 128 ± 23 in the Effort condition and 202 ± 51 in the Time condition. Subjects’ movement error rate was 40% (session #1: 46%; #2: 32%) in the Reference condition, 22% (session #1: 24%; #2: 19%) in the Effort condition and 48% (session #1: 49%; #2: 44%) in the Time condition. When movements were adequate, the overall percentage of correct choices was 80 % across the population (Reference condition: 79%; Effort condition: 83%; Time condition: 80%).

### Control analysis: effect of motor conditions on movement kinematic

To verify that the motor conditions effectively induced time- or energy-consuming reaching movements with respect to a reference condition, we analyzed reaching velocity peak and duration in each of the three motor blocks. We only report data collected in the choice task but results are similar in the DR task. Only trials in which an adequate movement was performed to express a choice, irrespective of the outcome of that choice, were included. As expected, reaching movement velocity peaks and durations were significantly modulated by the motor context in which movements were executed. Figure 2A shows each subject’s mean reaching velocity profile averaged across trials as a function of the motor condition. On average (± SD), for a similar duration (642 ± 85ms versus 597 ± 71ms), movement peak velocity was about twice higher in the Effort condition compared to the Reference (28.5 ± 5.3 cm/s versus 13.3 ± 2.6 cm/s, |z| = 246.0, p<0.001), whereas movement duration was about twice longer in the Time condition compared to the Reference condition (1076 ± 159ms versus 597 ± 71ms, |z| = 245.6, p<0.001). In the Time condition, the average (± SD) movement peak speed was slightly higher compared to the Reference condition (15.2 ± 3.3 cm/s versus 13.3 ± 2.6 cm/s, |z| = 32.1, p<0.001), but still much lower compared to the Effort condition (28.5 ± 5.3 cm/s, |z| = 213.3, p<0.001).

As noted above, it has been proposed that reaching effort is subjectively perceived as the temporally discounted metabolic cost of performing the movement. By estimating the effort of each reaching movement using equation 2, we observed that the effort level associated with executing reaching movements in the Effort condition is largely increased compared to the

Reference condition (−51.3 ± 13 J/s versus -34.4 ± 8 J/s, |z| = 165.0, p<0.001, Figure 2B). In the Time condition however, by imposing the same motion speed as in the Reference condition and doubling the distance to be covered, subjects’ reaching effort is much comparable, yet still slightly higher different, to the Reference condition (−35.2 ± 8 J/s versus -34.4 ± 8 J/s |z| = 8.3, p<0.001, Figure 2B).

### Effect of motor conditions on motor initiation

We first address the effects of motor conditions on action initiation. Indeed, the fully instructed delayed-reach (DR) task allows to assess the effects of the motor context on non-decision delays, mainly reflecting the motor initiation process. To do so, we compared subjects’ average reaction times (RTs) under each motor condition of the DR task. At the population level, we found that RTs in the DR task were significantly longer in the Time condition, i.e. when reaching duration was longer, compared to the Reference condition (475 ± 79ms in Time and 420 ± 53ms in Reference, |z| = 12.13, p < 0.001). This result is highly robust at the individual level, as the increase of a temporal motor cost extended RTs in the vast majority of subjects (21/31, p<0.05) compared to the Reference condition (Figure 3, left). This increase of RT was not observed in the Effort condition at the population level (414 ± 53ms). The effect of effort was also less pronounced and more variable at the individual level (Figure 3, right). Within-subject data indeed show that energy-consuming movements usually led to similar RTs compared to the Reference block (p<0.05 for 23/31 subjects), although some participants (7/31) reacted significantly faster (p<0.05).

### Effect of motor conditions on decision behavior

To investigate subjects’ decision behavior in the choice task, we analyzed their decision durations (DDs) and success probabilities (SPs) as a function of the motor context in which choices were reported. We first found at the population level that DDs were significantly shorter in the Time condition than in the Reference condition (1033 ± 334ms versus 1104 ± 321ms, |z| = 9.91, p<0.001). This observation is robust within subjects, as about half (14/31) of them made significantly faster decisions when the required movement duration was doubled (Figure 4A, left). Only five subjects showed the opposite pattern, i.e. a decrease of decision speed in the Time condition compared to the Reference condition. By contrast, we found no significant difference in DDs between the Reference and Effort conditions at the population level (1104 ± 321ms versus 1092 ± 333ms). This is because effort had a mixed influence on decision speed at the individual level (Figure 4A, right). 13 out of 31 participants did not adjust their DDs in the Effort condition compared to the Reference block, 8 were longer to decide in the Effort condition compared to the Reference condition, and 10 participants showed the opposite pattern.

**Figure 4:**
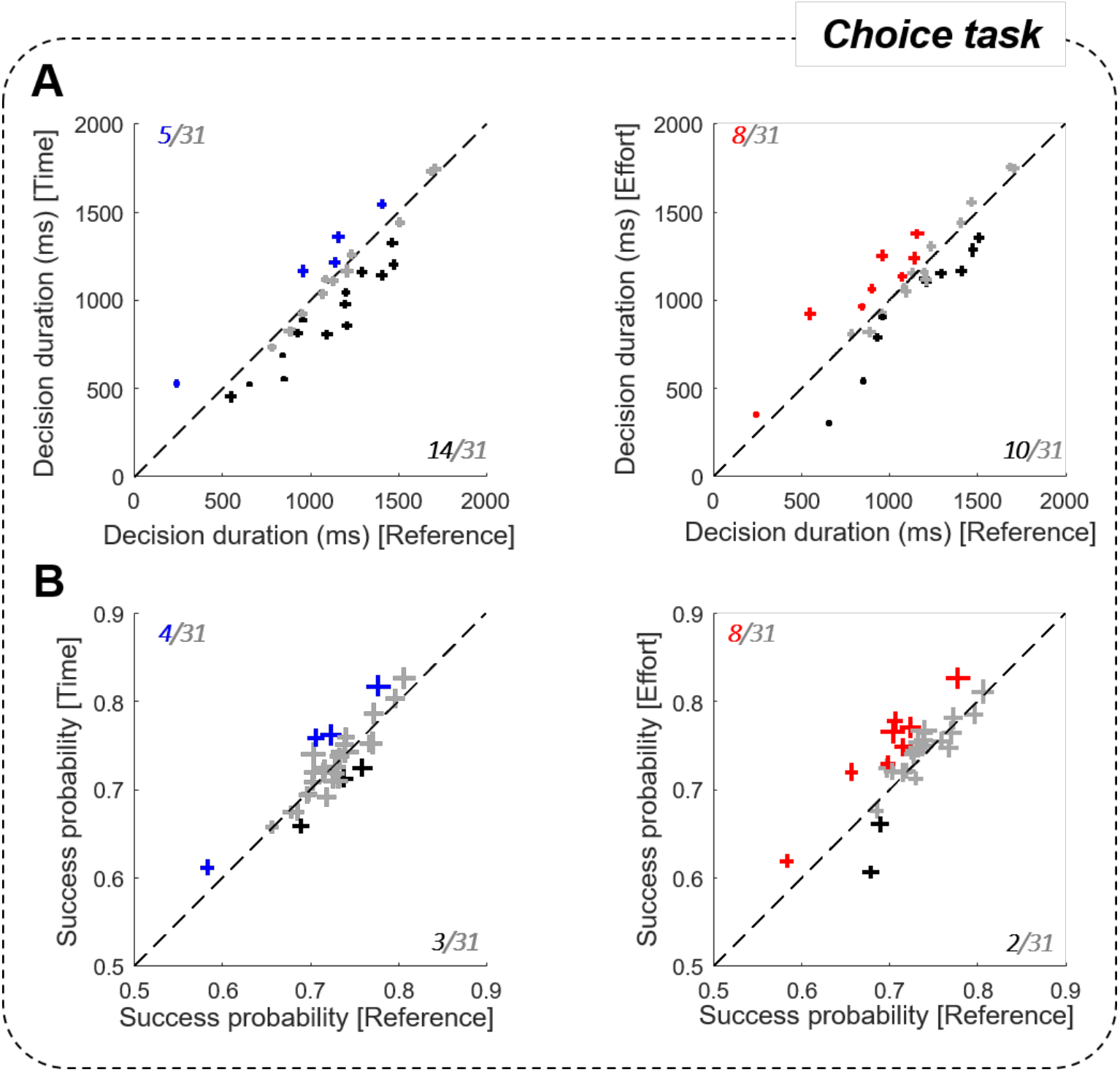
Effect of motor conditions on decision duration and success probability. **A**. Comparison of the mean (± SE) subjects’ decision durations between the costly conditions (left panel: Time / right: Effort) and the Reference condition in the choice task. Same conventions as in Figure 3. **B**. Same analysis as A for success probabilities at decision time.

To assess the consequences of a modulation of DDs on choice accuracy, we calculated subjects’ SPs at decision time (Figure 4B). At the population level, SPs at decision time were similar in the Time and the Reference conditions (0.73 ± 0.05 versus 0.72 ± 0.04) and were significantly higher in the Effort condition compared to the Reference condition (0.74 ± 0.05 versus 0.72 ± 0.04, |z| = 5.98, p<0.001). However, individual data revealed inconsistent effects, with only 8 subjects that showed a higher SP in the Effort condition compared to the Reference condition and for the majority of subjects (21/31), SPs were similar between conditions. Thus, despite the modulation of DDs described above, choice SP was not significantly impacted by movement duration and only marginally impacted by energy-consuming movements.

### Effect of motor conditions on the rate of reward

In each session of the choice task, subjects had to make 240 correct decisions, 80 in each motor condition. In each of these conditions, reaching movements carried a cost impacting subjects’ rate of reward in the current trial according to eq. 4. In this equation, the value (Δ, in Joules) assigned by the brain to a positive outcome in each trial is arbitrarily set to 500. We computed the expected rate of reward for each subject in each trial, averaged it across each motor condition trials and found at the population level that reward rate was lower in the Effort condition compared to the Reference condition (70 ± 6 versus 74 ± 6 J/s, |z| = 15.3, p < 0.001, Figure 5A). Reward rate was even more significantly reduced in the Time condition (60 J/s) compared to the Reference condition (63 ± 7 versus 74 ± 6 J/s, |z| = 40.7, p < 0.001, Figure 5A). These observations are robust at the individual level since all subjects had a lower reward rate in the Time condition compared to the Reference condition (p<0.05). In the Effort condition however, effects were more variable within subjects, as reward rate decreased for 20 participants, it increased for 1 subject and it did not significantly vary for the remaining 10 subjects compared to the Reference condition.

**Figure 5:**
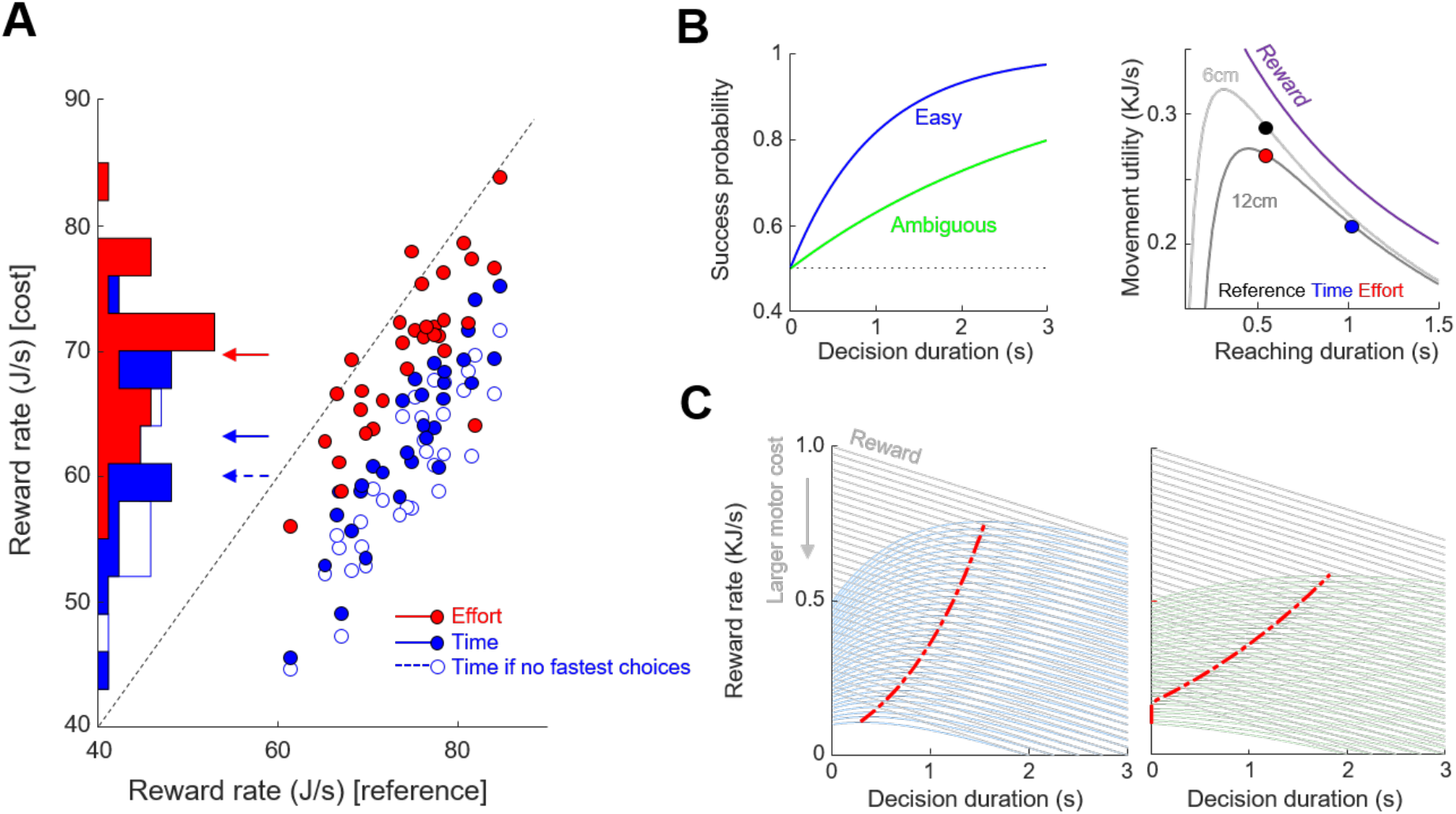
Observed and theoretical effects of motor conditions on reward rate. **A**. Distributions and comparisons of the average reward rate of each subject (dots) between costly conditions (ordinate, filled blue dots for Time, red for Effort) and the Reference condition (abscissa). Arrows mark the mean values for each condition. Unfilled blue dots show the rate of reward if subjects who shortened their decision durations in the Time condition compared to the Reference condition did not accomplish this shortening. **B**. Left: Hypothetic success probability (SP) profiles of one easy (blue) and one ambiguous (green) trials. Right: Reward temporal discounting and movement utility according to movement duration (abscissa) and amplitude (see text for details). **C**. Reward rate computed as the product between the reward value linearly discounted during decision duration (gray lines) and the SP (panel B, left) in easy (left panel) and ambiguous (right) trials for various initial reward values. The initial value of reward is reduced as motor cost increases (downward gray arrow). The red dots mark the maximal value for each reward rate function.

It is interesting to note that the most penalizing motor condition (the Time condition) is also the one in which most of the decision adjustments occurred, mostly in terms of decision duration. We describe in the following paragraphs a theoretical demonstration that offers an explanation linking these adjustments and the participant willingness to maximize their rate of reward. Suppose we consider a family of hypothetical success probability functions:

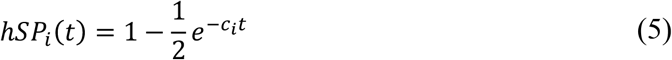

where *ci* is a parameter controlling trial difficulty that can vary from trial to trial. For the present demonstration, we simulate one hypothetic easy trial (Figure 5B, left, blue curve) and one hypothetic ambiguous trial (Figure 5B, left, green curve). Because these functions increase monotonically with time, waiting until the end of the sensory evidence presentation before committing sounds like the best policy in the choice task. However, spending time to collect sensory information also delays the acquisition of the reward, and time discounts the value of that reward. Assuming that in the choice task the reward is linearly (for simplicity) discounted by time *t* according to the function (Figure 5C, gray lines):

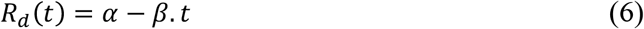

where *α* is the value assigned by the brain to a positive outcome given movement utility Um (see below), and *β* is the rate of discount, then we can define a theoretical reward rate function during the deliberation process as (Figure 5C, blue and green curves):

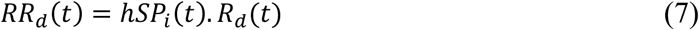

Because of the assumptions made above about *hSPi*(*t*), *RRd* has a single peak for each trial *i*. On any given trial, the probability of success starts at a point (0.5) and grows at some rate (fast in easy trials, more slowly in ambiguous trials). As long as the peak of the *RRd* function is not crossed, then one should continue to process information. However, as soon as that peak is crossed, one should commit.

Note that we include the cost of executing a movement in the definition of the rate of reward during the decision process (in *Rd*(*t*)). To quantify this cost, we assume that each movement carries a penalty, with respect to its duration and its energetic expenditure, that reduces the initial value (Δ) of the reward at the beginning of the trial. As mentioned in the introduction, movement duration and energetic expenditure are two costs that are intertwined in a trade-off: executing the slowest movements in order to minimize effort sounds like a good strategy, but passage of time *t* during movement discounts reward value, just like time discounts reward value during deliberation. For this demonstration, the temporal discounting of reward value during movement is expressed as (Figure 5B, right, purple curve):

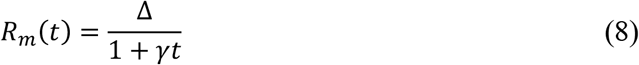

where γ =1 determines how rapidly reward is discounted (Shadmehr et al. 2016). As a result, the utility of the movement is expressed as the sum of the temporally discounted reward value and the temporally discounted energy consumption (or effort, eq. 2) during movement production:

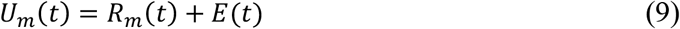

Thus, if we assume that the context-dependent utility of the movement impacts the rate of reward that subjects expect in a trial, theory predicts that in order to keep reward rate at its maximum when movement cost increases (or movement utility decreases), decision duration should be shortened, especially when the trial is difficult (see the red dots that mark the maximum value of the reward rate functions in Figure 5C).

In our experiment, movements executed in the Time condition carried much less utility (assuming the present Δ=500 and γ=1 parameters) compared to movements executed in the Reference and, to a lesser extent, in the Effort conditions (see the color dots in Fig. 5B, right).

Theory thus suggests that the 14 subjects who shortened their decision duration in the Time condition compared to the Reference condition did so to limit a drop of rate of reward induced by the strong temporal cost associated with executing reaching movements in this condition. If these subjects did not speed up their choices in the Time condition, their rate of reward would have been lower (60 versus 63 J/s, |z|=4.9, p<0.001, Figure 5A, open blue dots).

## DISCUSSION

Adapted behavior involves computations during which multiple trade-offs between reward value, accuracy requirement, energy expenditure and elapsing time need to be solved so as to obtain rewards as soon as possible while spending the least possible amount of energy. However, whether, how and why animals integrate movement time and energy costs into a decision-making policy is not fully understood. In this study, we asked 31 healthy human subjects to perform a perceptual decision-making task where the motor context in which a choice is reported was manipulated to dissociate the role of movement time and energy costs on participants’ decisions. We found that most subjects were influenced by motor costs during their deliberation process. Both duration and energy expenditure impacted decision-making but increasing reaching duration affected decision and motor initiation more consistently than increasing reaching energy expenditure. While time-consuming movements strongly extended reaction times in a fully instructed task compared to a reference condition, they often led to faster decisions in the choice task. We propose that subjects who shortened their choices in the time-consuming condition did so to limit a drop of reward rate at the session level. Importantly, effects of costs on decision-making and motor preparation often varied between subjects, especially when movement energy was manipulated, suggesting an idiosyncratic nature of the motor cost integration during goal-oriented behavior.

### Decision computations take motor costs into account

Decision-making has been traditionally described as a process that is completed prior to the preparation and execution of the action that reports the choice (Padoa-Schioppa 2011; Pylyshyn 1984). In ecological scenarios however, sensory or value-based decisions are very often expressed by actions that are themselves associated with risks and costs (Cisek and Kalaska 2010). In line with this embodied view of the decision process, the present results indicate that motor costs are part of the decision-making and movement initiation computations. Among costs, both duration and energy expenditure discount the value of rewards (Shadmehr et al. 2019; Shadmehr and Ahmed 2020). As a consequence, individuals tend to decide and act in a way that reduces these costs. For instance, when humans make rapid choices between reaching movements, they choose actions that carry the lowest biomechanical cost (Cos et al. 2011). Moreover, when the decision primarily relies on perceptual information, human subjects are biased in their decisions depending on the physical effort or the biomechanical cost associated with the movement executed to report a choice. If one movement carries a large cost, the probability of choosing that option decreases, even if the movement by itself does not influence success probability (Burk et al. 2014; Hagura et al. 2017; Marcos et al. 2015). In these studies, however, each of the two potential targets was assigned with a specific motor cost, and the relative contribution of movement energy expenditure and duration was not addressed. In the present work, the two targets were always associated with the same motor cost, and time and energy costs were independently varied between blocks of trials. This design allowed us to study the relative contribution of time- and energy-consuming motor contexts on subjects’ perceptual decision strategy. Recent studies addressing the relative contribution of motor costs on motor decisions suggest that time-related costs are the most impactful ones. Morel et al. (2017) found for instance that humans avoid time-consuming movements more often than other types of costly movements. Michalski et al. (2020) observed that movement amplitude, direction and accuracy influence the probability of switching from one ongoing movement to another more than energy expenditure. Our results support these studies by showing that varying movement duration impacts perceptual decision-making and movement initiation more often and more consistently than varying movement energy expenditure.

### Decision and action are two modes of one integrated process

Why would the motor context in which a decision is made have an influence on the decision itself? During natural behavior, decision and action are tightly linked. It is thus natural to imagine that both functions could share operating principles to maximize behavior utility. Indeed, for anyone making a decision, the most adaptive strategy is to choose options that maximize one’s global reward rate (Balci et al. 2011; Bogacz et al. 2010), which occurs when *both* decision and action are sufficiently accurate but not overly effortful and time consuming. In this view, decision and action define a continuum, coordinated by unified or interacting choice and motor regulation signals (Carland et al. 2019; Cisek and Thura 2018; Shadmehr et al. 2019; Thura and Cisek 2016, 2017). Recent observations support such coordination between decision and action during goal-directed behavior (Reynaud et al. 2020; Thura 2020; Thura et al. 2014; Yoon et al. 2018).

In the present delayed reaching task in which both where and when to reach were instructed, the imposed extended movement duration increased reaction times for the vast majority of the subjects. Usually, if the distance to a rewarded target increases, individuals increase their reaching speed to limit the impact of the temporal discounting of reward (Reppert et al. 2018). In the present study however, subjects could not reduce movement duration. It is thus possible that the larger temporal discounting of reward expected by subjects in this context reduced their implicit motivation to behave (Mazzoni et al. 2007; Shadmehr et al. 2019), leading to longer reaction times. By contrast, almost half of the subjects reduced their decision durations in the Time condition of the choice task compared to a control condition. Because the Time condition strongly reduced subjects’ expected reward rate (Figure 5A), it is possible that those subjects attempted to compensate the time-consuming movements by reducing their choice duration during the deliberation period. Theoretical simulations (Figure 5B) indicate that such strategy limits a drop of reward rate. This observation suggests a flexible mechanism allowing to trade decision speed for movement speed in order to maintain a decent rate of reward despite constraining motor conditions (Reynaud et al. 2020). Interestingly, the analysis of error movement trials (too short or too long with respect to the instructed temporal interval, see Methods) supports such an integrated view of goal-directed behavior. Figure 6 shows that the too short movements were overall made when decisions were long, and the too long movements were made when decisions were short (1150 ± 59ms versus 1042 ± 57ms, |z|=12.5, p<0.001), suggesting that many subjects (21/31, p<0.05) were primarily concerned about computing a global trial duration rather than computing decision and action durations separately.

**Figure 6:**
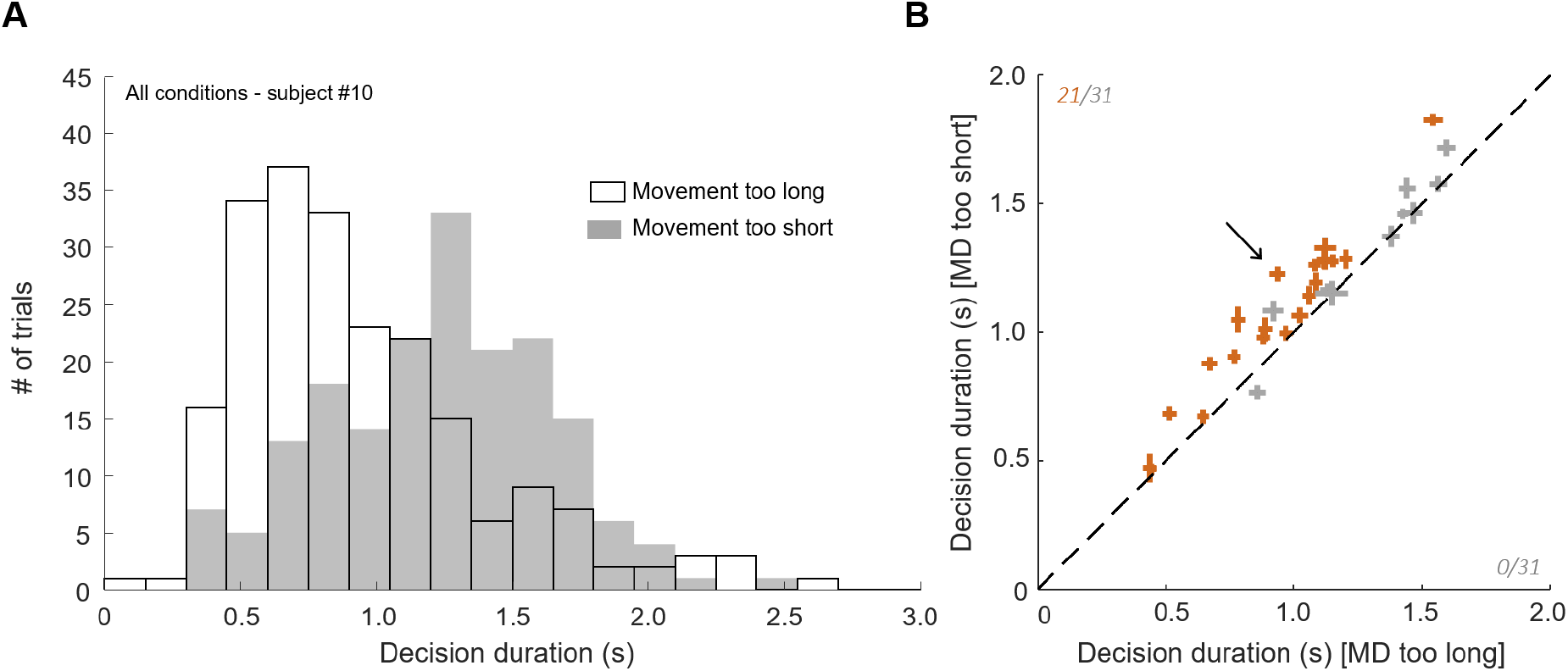
Decision duration in error movement trials. **A**. Distribution of one example subject’s decision durations with trials sorted according to the reaching error type, i.e. too long or too short. Trials performed in the three motor conditions are included. **B**. Mean ± SE decision duration for each subject in trials during which movements were either too long (abscissa) or too short (ordinate). The arrow indicates the subject shown in A. Same convention as in Figures 3 and 4.

### The question of the reaching effort cost

Compared to the effects of the time-consuming movements discussed above, the impact of the energy-consuming movements on decision and action initiation were less pronounced, especially in the DR task, and more variable at the population level. This result does not fully support the implicit motor motivation hypothesis (Mazzoni et al. 2007), according to which effortful movements discount reward value, thus motivation, delaying movement initiation and reducing movement vigor (Shadmehr et al. 2019; Summerside et al. 2018; Wickler et al. 2000). We even observed the opposite effect for 7 and 10 out of 31 subjects in the delayed reaching and the choice tasks, respectively. For those subjects, it is possible that the instruction to produce more vigorous movements energized their behavior at a global level, leading to faster choices and shorter movement initiations as predicted by the shared regulation hypothesis (Carland et al. 2019; Cisek and Thura 2018; Thura 2020; Thura et al. 2014). By contrast, subjects who spent more time to make their decisions in the effortful condition compared to the control condition possibly aimed at collecting more sensory evidence to avoid choice errors and to ultimately minimize the total number of trials to perform. Alternatively, the more metabolically demanding movements may have, as mentioned above, diminished subjects’ motivation to perform the task, leading to longer decisions (Mazzoni et al. 2007).

The lack of consistent effects of reaching effort on decision making and movement initiation at the population level can be explained by several reasons. First, movement effort could not be as directly compensated in the choice task as the time-related cost. Indeed, the deliberation period of the choice task gives a very large window for temporal adjustments, whereas the task does not incorporate a similar effort domain that could have allowed energy costs to be as directly compensated. Moreover, in the present study, subjects faced only two levels of motor effort. It is thus possible that the chosen parameters felt outside of the range that would have been efficient to affect subjects’ behavior in a more consistent manner. Finally, the way to manipulate movement energy expenditure often differs between studies. It can be performed through variation of movement trajectories, leading to different biomechanical costs (Cos et al. 2011), through loads or resistances applied on the moving segments (Morel et al. 2017), through isometric manipulations such as handle squeezes (Körding et al. 2004), etc. Here we chose to manipulate movement energy expenditure by imposing various durations for a given amplitude, thus manipulating movement speed. The relation between movement speed and energy expenditure has been well documented, and it offers a convenient theoretical framework to study the impact of motor costs on decision-making (Shadmehr et al. 2016; Shadmehr and Ahmed 2020). It is still possible that some of the present results would have been different if other types of effort manipulations had been performed.

### Influence of motor costs across subjects and behavioral repertoires

The present results also indicate that motor costs effects on decision and action initiation are variable at the population level, especially with respect to energy expenditure. This observation was expected as previous work demonstrated that motor cost is a subjective estimation that does not impact behavior in a consistent way across individuals. For instance, some people consider the effort produced during physical activities as a reward whereas others tend to favor sedentary behavior (Cheval et al. 2018b, 2018a). Regarding elapsing time, Choi and colleagues (2014) showed in a saccadic task that some individuals exhibit a much greater sensitivity to temporal costs than others. Similarly, Berret and colleagues (2018) showed that self-selected vigor of pointing movements strongly differs between subjects but tend to be relatively constant within each subject, even when biomechanics-related costs are taken into account. These results suggest that both effort sensitivity and the temporal discounting rate differs between people, possibly explaining why some subjects did not adjust their decision policy as a function of motor costs in the present study while other did. In addition, the temporal discounting rate varies throughout individual’s life as well. In humans, the temporal discounting tends to be steepest in adolescence and then declines with age (Green et al. 1999). Despite the present population age range was rather narrow, we cannot exclude that age influenced how some subjects reacted with respect to motor costs in both tasks.

Finally, it’s worth mentioning that subjects expressed their choices via reaching movements. One may also ask whether similar effects would have been observed in other movement types, such as locomotion or saccades. Locomotion-related computations seem to share many of the utility principles described for reaching movements. For instance, the preferred walking speed correlates with a minimization of the metabolic cost during displacement (Donelan et al. 2001; Summerside et al. 2018; Zarrugh et al. 1974). Moreover, consistent inter-individual differences of vigor between reaching and walking tasks are generally found (Labaune et al. 2020). By contrast, the oculomotor system seems to differ from the reaching and the locomotion systems with respect to behavior utility (Labaune et al. 2020; Reppert et al. 2018). More studies are needed to test the questions addressed in the present work in other behavioral repertoires.

## ACKNOWLEDGMENT

This work was supported by a CNRS/Inserm ATIP/Avenir grant and an Inserm young investigator fellowship to DT. The authors thank Sonia Alouche and Jean-Louis Borach for effective administrative assistance, Frédéric Volland for his expertise during the technical preparation preceding this experiment, David Robbe and Bastien Berret for their contributions to the development of the ideas addressed in this manuscript.

